# Perceptual coupling and decoupling of the default mode network during mind-wandering and reading

**DOI:** 10.1101/2020.10.03.324947

**Authors:** Meichao Zhang, Boris Bernhardt, Xiuyi Wang, Dominika Varga, Katya Krieger-Redwood, Jessica Royer, Raul Rodriguez-Cruces, Reinder Vos de Wael, Daniel S. Margulies, Jonathan Smallwood, Elizabeth Jefferies

## Abstract

While reading, the mind can wander to unrelated autobiographical information, creating a perceptually-decoupled state detrimental to narrative comprehension. To understand how this mind-wandering state emerges, we asked whether retrieving autobiographical content necessitates functional disengagement from visual input. In Experiment 1, brain activity was recorded using functional magnetic resonance imaging (fMRI) in an experimental situation mimicking naturally occurring mind-wandering, allowing us to precisely delineate neural regions involved in memory and reading. Individuals read expository texts and ignored personally relevant autobiographical memories, as well as the opposite situation. Medial regions of the default mode network (DMN) were recruited during memory retrieval. In contrast, left temporal and lateral prefrontal regions of the DMN, as well as ventral visual cortex, were recruited when reading for comprehension. Experiment 2 used functional connectivity at rest to establish that (i) DMN regions linked to memory are more functionally decoupled from regions of ventral visual cortex than regions in the same network engaged when reading, and (ii) individuals reporting more mind-wandering and worse comprehension, while reading in the lab, showed increased functional decoupling between visually-connected DMN sites important for reading and a region of dorsal occipital cortex linked to autobiographical memory in Experiment 1. These data suggest we lose track of the narrative when our mind wanders because the generation of autobiographical mental content relies on cortical regions within the DMN which are functionally decoupled from ventral visual regions engaged during reading.

**Significance statement:** When the mind wanders during reading, we lose track of information from the narrative. We hypothesised that poor comprehension occurs because retrieving autobiographical memories reduces the perceptual coupling necessary to understand written words. We show that default mode network (DMN) areas involved in reading are functionally more connected to ventral visual regions than DMN regions important for autobiographical memory. Furthermore, individuals who mind-wander more, and comprehend less, have weaker connectivity between visually-coupled DMN regions linked to reading and dorsal occipital areas linked to autobiographical memory. These data suggest that when our minds wander during reading, retrieval of personally-relevant information activates DMN regions that are functionally disconnected from visual input, creating a perceptually decoupled state detrimental to comprehension.

## Introduction

The human mind is remarkably flexible, capable of shifting focus from information in the external environment to perceptually-decoupled states that are generated from information in memory (1–3). This capacity for self-generating mental content is ubiquitous across cultures and has links to both beneficial and detrimental features of cognition (4). Although self-generated states are common in daily life (5, 6), they can be problematic if they occur during reading (7, 8). The detrimental effects of mind-wandering on reading are believed to occur because this state elicits perceptual decoupling that disrupts narrative comprehension (1, 9).

Contemporary work in cognitive neuroscience has shown that both reading for comprehension (10), and off-task states (11, 12), are linked to activity within the default mode network (DMN). For example, the building blocks of reading for comprehension are conceptual representations supported by regions overlapping with DMN in the anterior, ventral and lateral temporal lobe (13). In contrast, the content of self-generated thoughts often comes from autobiographical memory (14), linked to medial regions of the DMN including posterior cingulate, ventral prefrontal and inferior parietal cortex (15). Furthermore, studies of individual differences highlight that both better reading comprehension, as well as greater tendency for off-task thought, are predictable based on neural patterns in the DMN, as well as in other cortical regions (16, 17). For example, individuals who are better at reading for comprehension show more functional integration between lateral and medial elements of the DMN, while individuals who tend to be more off-task show greater decoupling between the DMN and regions of visual cortex important for visual processing during reading (16, 17).

Converging empirical and theoretical evidence, therefore, suggests that both reading for comprehension, and off-task self-generated states, depend on regions within the broader DMN. Recent views suggest the DMN’s role in cognition emerges from this system’s topographic location, with core nodes located in regions that are distant in both functional and structural terms from unimodal cortex (18). This topographical organisation has been argued to explain the role of the DMN in multiple cognitive states because it locates this network at the end of information processing streams that are necessary for relatively abstract tasks (like reading comprehension) but also explains why the same network can be involved in states that require disengagement from the here and now (such as mind-wandering; 19). This topographically-informed view of the DMN provides a novel hypothesis for why mind-wandering during reading creates a situation in which we lose track of the meaning of the words we are reading. We anticipate that the process of generating mental content using information from memory leads to a perceptually-decoupled state associated with poor comprehension. When this occurs, there is a shift in the balance of neural activity within the DMN, away from DMN regions functionally coupled with ventral visual regions important for reading comprehension, to other regions within DMN that are more isolated from inputs.

To test this account of the consequences of mind-wandering while reading, we conducted two experiments using functional Magnetic Resonance Imaging (fMRI) to measure brain activity. In Experiment 1, participants (N = 29) performed tasks that mimicked the experience of mind-wandering while reading. In one condition, participants were presented with information from an expository text on the screen but asked to scan these words while instead retrieving a personally relevant autobiographical memory. In a second condition, participants focused on a similar expository text, while refraining from attending to autobiographical information. Our experiment exploits the fact that self-generated states can be understood as the spontaneous engagement of processes that can also be recruited as part of a task (1). This process view allowed us to identify, with a high degree of precision, the neural regions involved in the two cognitive components of interest: autobiographical memory retrieval and reading for comprehension. Experiment 2 examined the functional architecture of regions involved in these two states, evaluating whether the DMN regions important for reading are more functionally connected to the ventral visual stream than the DMN regions involved in autobiographical memories (N = 243). Finally, we also sought to generalise these results by examining whether functional relationships between regions implicated in our experimental task, which was designed to mimic mind-wandering while reading, could predict individual differences in naturally-occurring mind-wandering during reading (N = 69).

Foreshadowing our results, we found that (a) left lateral prefrontal and temporal regions of the DMN, within the dorsomedial subsystem (20, 21), are important for reading, while regions of the core DMN are engaged during autobiographical memory (medial prefrontal, posterior cingulate and angular gyrus); (b) regions of the DMN linked to reading are more functionally connected to ventral visual regions than DMN regions implicated in autobiographical memory, and (c) DMN regions linked to reading are decoupled from a region in dorsal occipital cortex linked to autobiographical memory for individuals who naturally mind-wander more during reading, generalising our experimental paradigm to an ecologically valid situation. Together these data support the hypothesis that processes involved in the generation of mental content from memory depend on regions in the DMN that are functionally decoupled from ventral visual regions important for reading. Our analysis, therefore, suggests that the reason why we lose track of the narrative when our mind wanders during reading is because the generation of autobiographical content relies on neural activity in core DMN regions and this encourage a state of perceptual decoupling that reduces comprehension of external input.

## Results

### Experiment 1

#### Behavioural results

Our first goal was to establish whether our experimental situation successfully mimicked features of mind-wandering while reading, namely (i) a focus on personally-relevant information accompanied by (ii) a reduced focus on the text. A repeated-measures Analysis of Variance (ANOVA) was used to examine this question by comparing the effects of autobiographical memory retrieval on reading (and vice versa; see the left hand of Figure 1). Participants rated their task focus for each trial on a scale of 1 (i.e. not at all) to 7 (i.e. very much) in the scanner and reported reduced task focus on the primary task for both reading and autobiographical memory retrieval when both tasks were presented at the same time (i.e. in Conflict conditions), *F*(1,28) = 44.28, *p* < .001, *η_p_^2^* = .61 (see panel A in the right hand of Figure 1). There was no interaction between Task and Conflict, *F*(1,28) = .12, *p* = .73, *η_p_^2^* = .004, indicating that the effect of conflict between tasks had an equivalent effect on mental focus in reading and autobiographical memory. For the reading trials, retrieval of autobiographical memories reduced rated comprehension, *F*(1,28) = 10.40, *p* = .003, *η_p_^2^* = .27, but not participants’ level of prior familiarity with the sentence material, *F*(1,28) = 2.10, *p* = .16, *η_p_^2^* = .07 (see panel B in the right hand of Figure 1). For autobiographical memory, concurrent presentation of meaningful text reduced the vividness of the memories that were retrieved, *F*(1,28) = 36.29, *p* < .001, *η_p_^2^* = .56, as well as the rated consistency between retrieval in the scanner and the memory described for the cue word outside the scanner, *F*(1,28) = 8.19, *p* = .008, *η_p_^2^* = .23 (see panel C in the right hand of Figure 1). This pattern establishes that our paradigm successfully captures important features of the mind-wandering reading state, particularly the mutual inhibition between self-generated mental content and effective reading for comprehension.

**Figure 1.**
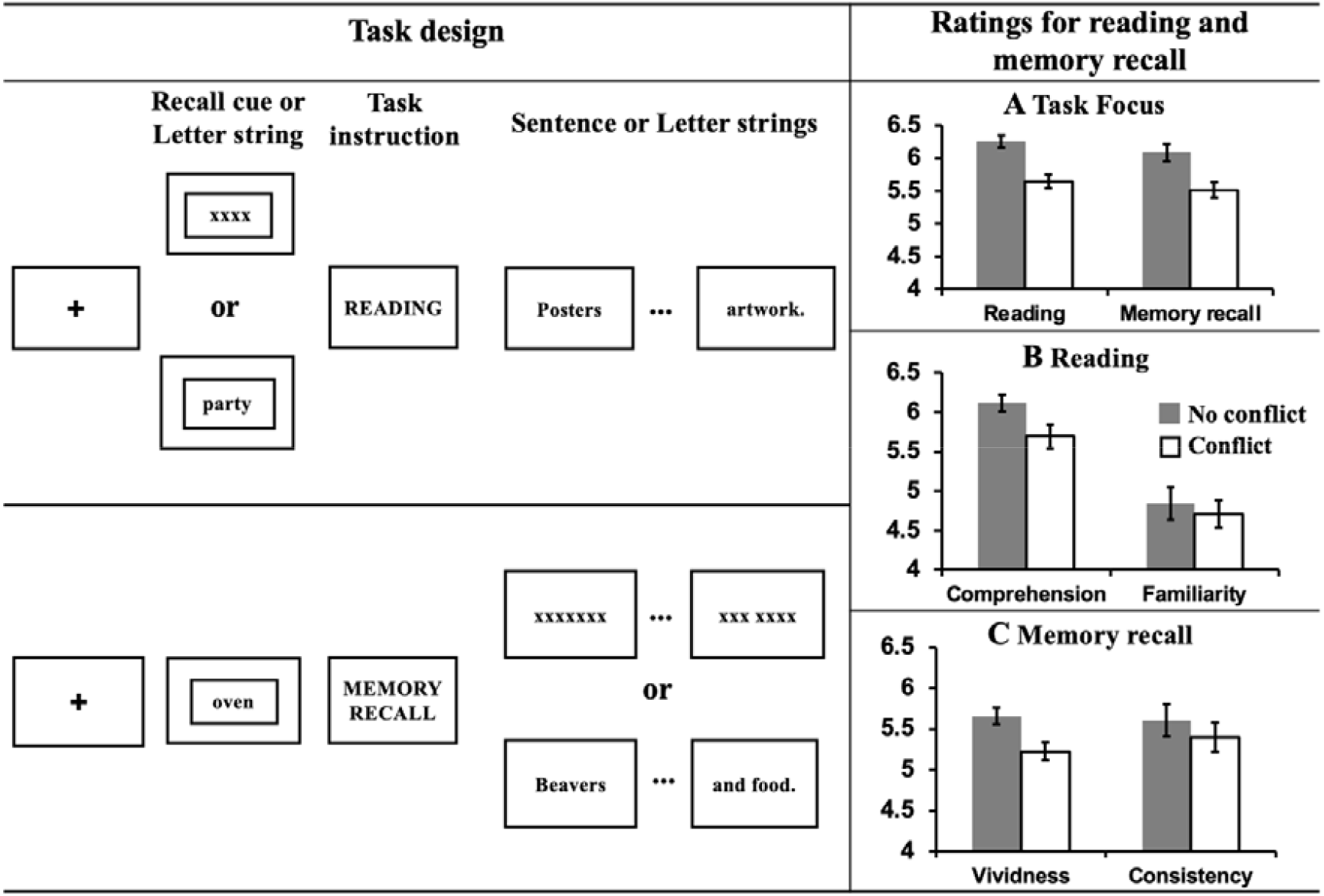
**Left hand panel: Task design – Experiment 1.** Using a counterbalanced design, participants either engaged in normal reading, or instead were focused on personally relevant information from memory. To mimic the mind-wandering while reading state, task conflict was created by presenting sentences during memory recall, or memory cues before the presentation of sentences. To understand the effect of meaningful input on memory retrieval, on some occasions ‘X’s were presented instead of text. To understand the effect of memory retrieval on reading, sometimes no memory was cued at the start of the reading trial. ***Right hand panel: Evidence of mutual inhibition between reading and autobiographical memory retrieval.* (A)** Participants rated their task focus as lower when reading while retrieving autobiographical memories (as well as vice versa). **(B)** Participants rated their comprehension of written material as lower when also retrieving autobiographical information. There was no effect on participants’ ratings of their familiarity with the content of the sentences. **(C)** Participants rated their autobiographical memories as less vivid and less consistent with their previously generated memories when meaningful text was presented at the same time. Error bars show standard error of the mean (SEM).

#### Neuroimaging results

Having established the expected pattern of competition between autobiographical memory retrieval and reading, we next considered the neural correlates that distinguish these states (Figure 2A). Contrasting blocks in which the primary task was reading with autobiographical memory retrieval highlighted a set of left lateralised regions within the temporal lobe and prefrontal cortex, including inferior frontal gyrus and superior and middle temporal gyri, which are recruited during reading. Activation in bilateral ventral visual cortex was also observed. In contrast, periods when autobiographical memory was the primary task were associated with greater neural activity in regions including medial and lateral prefrontal cortex, posterior cingulate cortex and angular gyrus. In Figure 2A, regions showing greater activity during reading are presented in warmer colours, and regions showing greater activity during autobiographical memory retrieval are presented in cooler colours.

**Figure 2.**
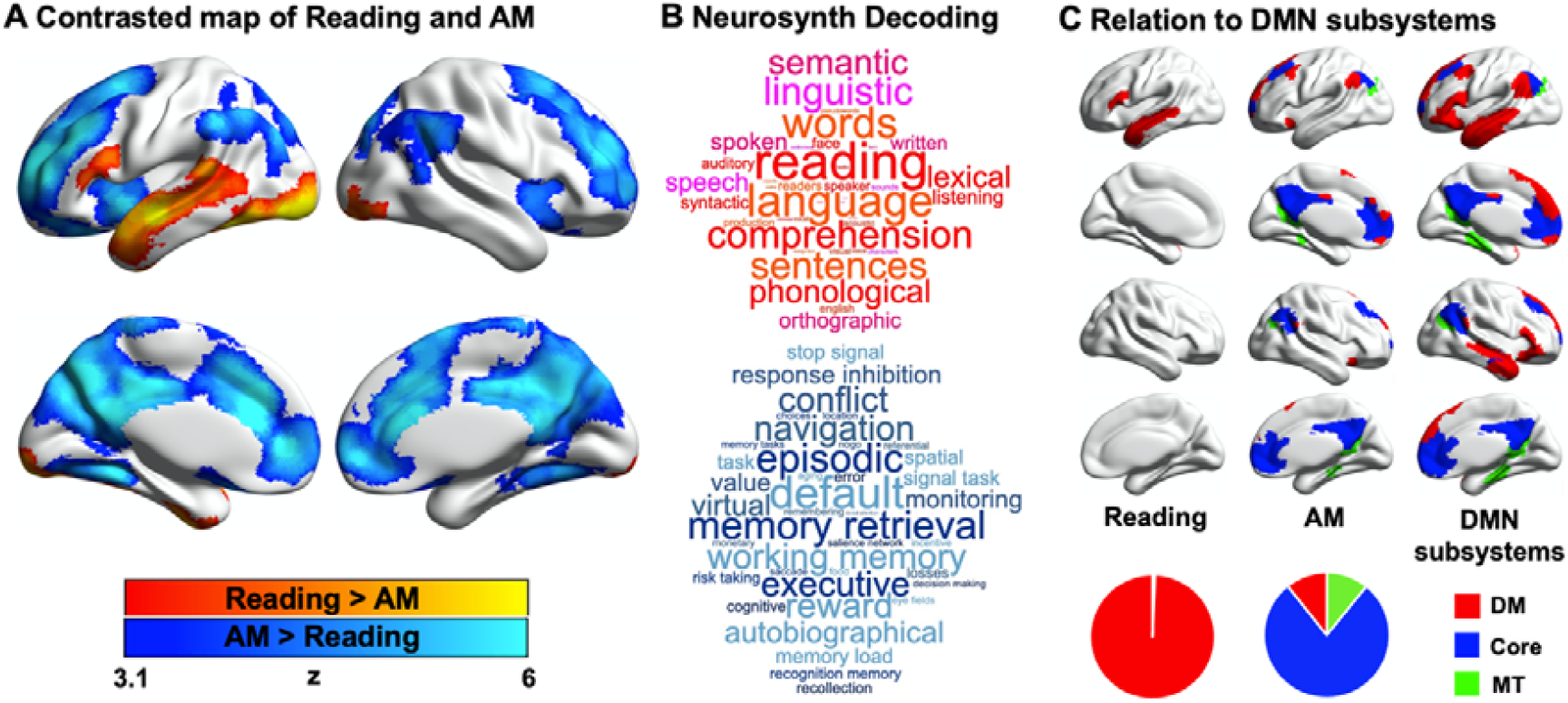
Neural activity associated with reading and autobiographical memory retrieval. **(A)** A comparison of regions showing significantly greater activity during reading (red) or autobiographical memory retrieval (blue). **(B)** A meta-analysis of the regions showing activity during reading and autobiographical memory retrieval using Neurosynth. In these word clouds the font size of the item illustrates its importance and the colour indicates its association (red = reading, blue = autobiographical memory). **(C)** Relationship between the patterns of observed activity during reading and autobiographical memory retrieval and their relationship to the subsystems of the DMN as described by Yeo and his colleagues (20). In this panel, regions in red fall within the dorsomedial (DM) subsystem, regions in blue fall within the core subsystem and regions in green fall within the medial temporal (MT) subsystem. The pie charts show the proportion of significant voxels associated with each condition that fall within each subsystem.

To confirm the most likely functional associations with these regions, we performed a metaanalysis using Neurosynth (see Methods). The results of this analysis are presented in Figure 2B in the form of word clouds where the font size describes the strength of the relationship and the colour describes the associated state (Red = reading, Blue = autobiographical memory retrieval). As expected, there was a correspondence between the psychological features of our conditions and the functional terms revealed by the meta-analysis, with regions linked to reading associated with terms such as “reading” and “language”, while regions linked to autobiographical memory retrieval were associated with terms like “memory retrieval” and “episodic”.

Prior studies have linked both semantic and autobiographical memory processes to the broader DMN, and we examined how the neural patterns associated with our states reflected the activation of the different subsystems of the DMN (20, 21). For each of the significant DMN voxels associated with reading or autobiographical memory retrieval, we examined whether they fell within the dorsomedial, core or medial temporal DMN subsystems, as defined by Yeo, Krienen and colleagues (20). The results of this analysis are presented in Figure 2C where the different columns show the different states (reading and autobiographical memory retrieval), and the different colours correspond to the DMN subsystems. The percentages of voxels falling within each system are presented as pie charts at the foot of this panel. It can be seen that DMN regions engaged during reading were entirely within the dorsomedial system (red; 100%), while the majority of the DMN voxels showing higher activity during autobiographical memory retrieval fell within the core subsystem (blue; 78%), with equal percentages in the dorsomedial (red; 11%) and medial temporal subsystems (green; 11%).

Having established that autobiographical memory and reading activate distinct subsystems within the broader DMN (despite some overlap in brain activation elicited by these states relative to the letter string baseline, see Figure S2), we next explored the functional consequences of neural activity in these conditions. Our behavioural analysis demonstrated a pattern of mutual competition between the two conditions (Figure 1); we therefore examined the relationship between the observed pattern of neural activity in each condition and a persons’ reported focus on the primary task. The results of this analysis are presented in Figure 3A where it can be seen that regions in medial prefrontal and parietal cortex, superior frontal gyrus and left lateral parietal cortex showed a stronger effect of task focus for autobiographical memory retrieval, relative to reading. We conducted a formal conjunction to identify how these regions mapped onto those showing differential activity for the two states. This is presented in Figure 3B, which shows that these clusters linked to better task focus for autobiographical memory also showed stronger activity during memory retrieval and lower activity during reading.

**Figure 3.**
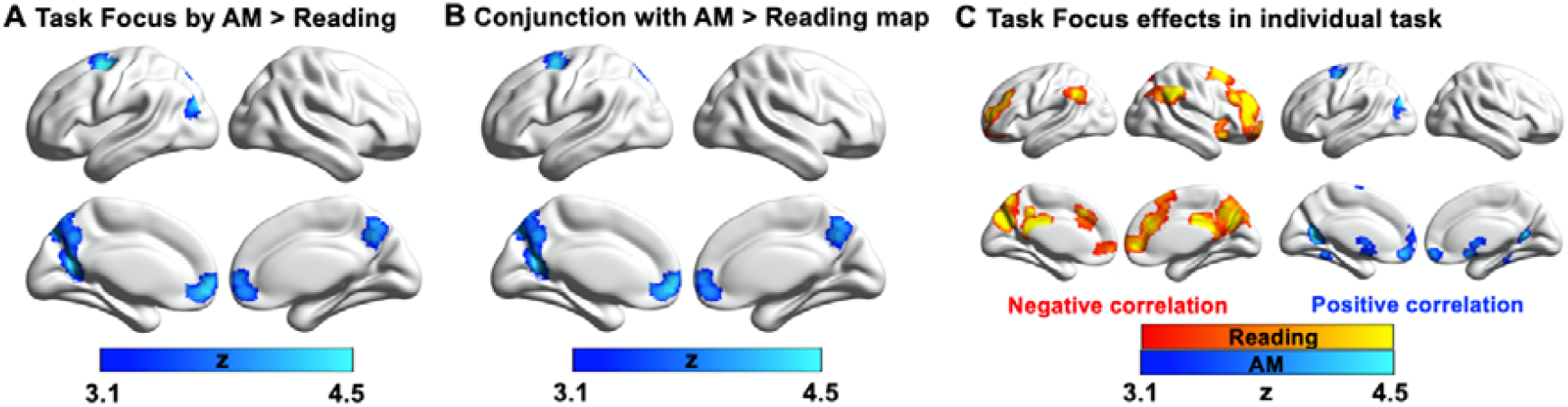
Parametric effects of task focus. **(A)** Regions showing a differential relationship with task focus across the two states (autobiographical memory and reading). These regions show greater activity when participants reported better focus on the task during autobiographical memory retrieval and poorer task focus during reading. **(B)** A formal conjunction between these regions with those showing greater activity across the tasks (Figure 2A). **(C)** Task focus effects in reading (red; negative correlation with task focus) and autobiographical memory recall task (blue; positive correlation with task focus).

Importantly, we confirmed that these regions showed significant associations with task focus separately in each task (see Figure 3C). Greater task focus during memory retrieval was associated with increased activation in medial prefrontal cortex, posterior parietal cortex (bordering dorsal lateral occipital cortex), superior frontal gyrus, retrosplenial cortex, and temporal fusiform cortex, including many sites within medial temporal and core DMN (50% and 40% of DMN voxels, respectively). In contrast, greater task focus in reading was correlated with increased deactivation in similar regions in bilateral middle/medial frontal gyrus, frontal pole, insular cortex, anterior/posterior cingulate gyrus, precuneus and angular gyrus – again, with substantial overlap with core DMN (82% of DMN voxels). This analysis establishes that regions of ventromedial prefrontal cortex, posterior cingulate cortex and superior frontal gyrus contribute to better focus on autobiographical memory retrieval while compromising the ability to read for comprehension. Greater task focus on autobiographical memory recall is linked to increased activation in core DMN regions, while deactivation of similar core DMN regions is linked to greater mental focus during reading.

### Experiment 2

Experiment 1 established that our paradigm captured the expected mutual inhibition between reading and autobiographical memory retrieval (Figure 1) and shows that both states depend on activity within distinct regions within the DMN (Figure 2 and 3). The aim of Experiment 2 was to understand (i) whether the DMN regions associated with reading and autobiographical memory retrieval show differences in their functional connectivity to ventral visual regions important for reading, and (ii) how individual differences in the connectivity of these DMN subsystems relate to the tendency to mind-wander during reading in a more naturalistic setting.

Our first analysis examined the extent to which DMN subnetworks are functionally connected to regions of the ventral visual stream activated during reading. This analysis helps us to determine whether the reductions in ventral visual cortex seen in Experiment 1 during autobiographical memory solely reflect an attentional phenomenon that emerges because participants were asked to attend to memory retrieval rather than textual input, or whether these effects relate to differences in the intrinsic functional architecture of the DMN regions important for autobiographical memory retrieval and reading. To address this question, we conducted whole-brain resting-state functional connectivity analyses targeting the regions of the DMN differentially associated with reading and autobiographical memory (see Methods). The results of this analysis can be seen in Figure 4A; regions more strongly correlated with DMN regions linked to reading are presented in warm colours, while those regions showing greater functional connectivity with DMN regions linked to autobiographical memory retrieval are shown in cool colours. It can be seen that DMN regions linked to reading show greater intrinsic connectivity to ventral visual cortex. We conducted a formal conjunction analysis between the regions showing stronger intrinsic connectivity with reading versus autobiographical memory activation sites within DMN and the spatial map of stronger activity during reading versus autobiographical memory retrieval. The results of this analysis are presented in Figure 4B. This “task-rest” conjunction establishes that the functional connectivity of regions linked to reading is mirrored by their joint activation during reading. Co-activation of aspects of DMN and visual cortex during reading is at least partially rooted in their intrinsic functional organisation. A supplementary analysis also revealed patterns of structural connectivity supporting this connection between ventral visual regions and areas of DMN that support reading (see Supplementary Figure S6).

**Figure 4.**
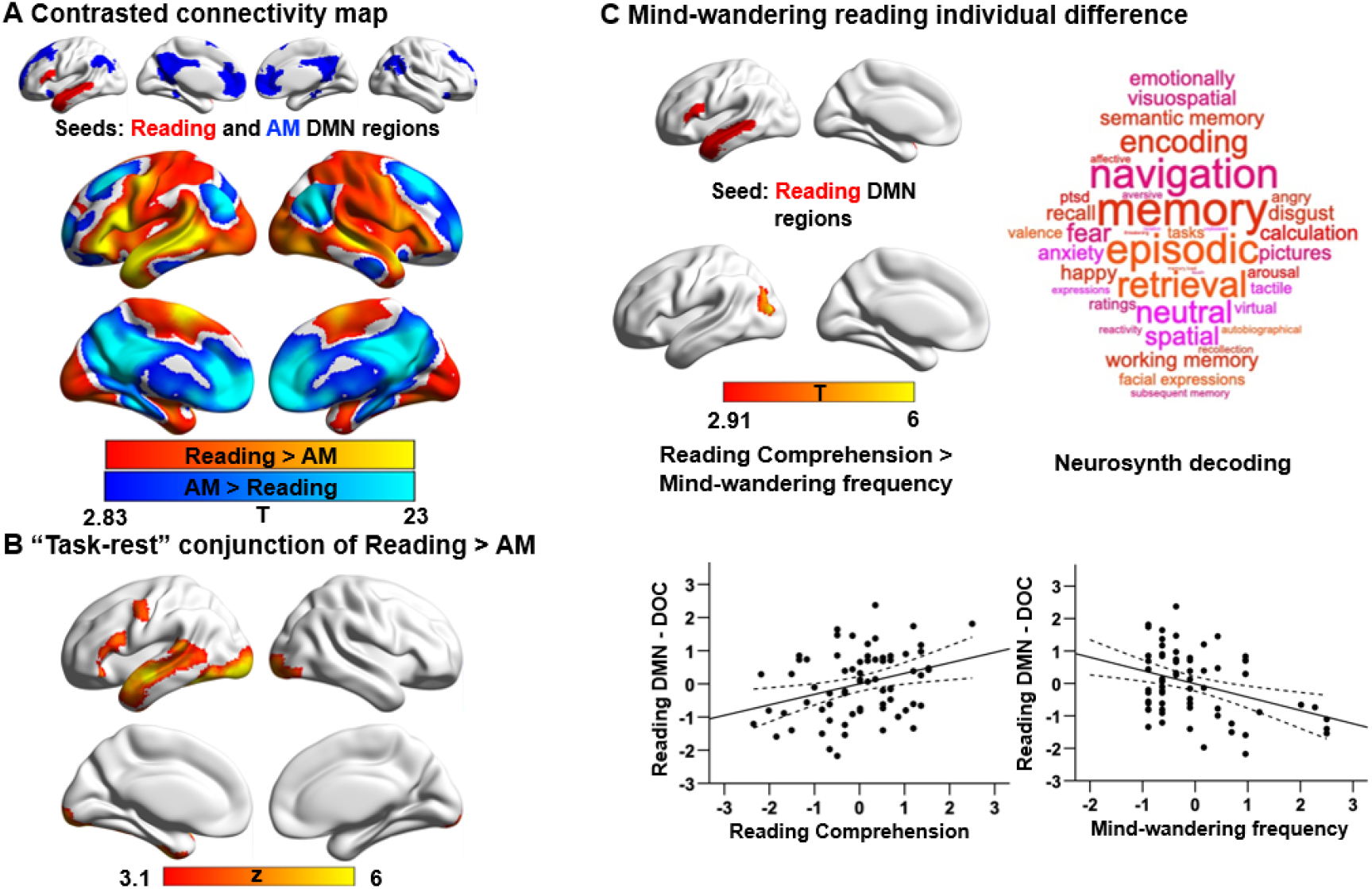
Results of Experiment 2: Analyses examining the functional architecture of DMN regions associated with reading and autobiographical memory and their relation to individual differences in mind-wandering reading. **Panel (A)** shows the results of a functional connectivity analysis showing differences between the DMN regions showing greater activity during reading (warmer colours) and autobiographical memory retrieval (cooler colours). The lower **panel (B)** shows results of a formal conjunction between regions associated with greater activity during reading versus autobiographical memory retrieval, and regions showing stronger correlation at rest with DMN regions more activated by reading. **Panel (C)** shows the relationship of these regions’ functional architecture and self-reports of mind-wandering reading. Group-level regression, using DMN regions showing greater activity during reading as a seed, demonstrated stronger connectivity with a region of dorsal occipital cortex (DOC) in individuals with good comprehension and infrequent mind-wandering. The scatterplots present the correlation between behaviour (reading comprehension: *r* = .32, *p* = .008, and mind-wandering frequency: *r* = .37, *p* = .002) and the average correlation with the reading-relevant DMN seed and the identified cluster (beta values). The error lines on the scatterplot indicate the 95% confidence estimates of the mean. Each point describes an individual participant. No comparable associations were observed for DMN regions showing greater activity during the retrieval of autobiographical memories. The word cloud shows the functional associations with this region of cortex performed using Neurosynth.

Our second analysis examined how the functional architecture of the DMN regions associated with reading and autobiographical memory retrieval relates to individual variation in naturally occurring mind-wandering during reading as measured in our prior study (17). We performed a group-level regression using the regions of the DMN linked to reading and autobiographical memories as seeds, and individual differences in reading comprehension and off-task thoughts (i.e., frequency of mind-wandering and the content of off-task thoughts including autobiographical memory; see Supplementary Figure S5) as explanatory variables. This revealed a region of dorsal occipital cortex bordering angular gyrus that showed less connectivity with regions of the DMN activated during reading for individuals who showed more frequent mind-wandering and poorer comprehension (see Figure 4C). This region largely overlapped with areas associated with task activation (AM > Reading: 52%; no overlap with Reading > AM map) and intrinsic connectivity (AM > Reading: 81%; Reading > AM: 8%) for autobiographical memory as opposed to reading, in Figure 2A and 4A. To confirm the most likely functional associations with this site we performed a meta-analysis using Neurosynth. The results of this analysis are presented in Figure 4C in the form of word clouds where the font size describes the strength of the relationship. This decoding of this region yielded terms largely linked to memory, such as “episodic” and “retrieval”. Finally, in a supplementary analysis, we examined the consequences of this variation in connectivity for the brain’s functional architecture (Supplementary Figure S7). Participants from an independent sample who had stronger intrinsic connectivity between reading-related DMN regions and dorsal occipital cortex (i.e. the pattern of connectivity associated with better reading comprehension and less mind-wandering) also showed stronger connectivity between reading-related visual regions and the same site in dorsal occipital cortex. In summary, this study establishes a correspondence between our experimental context that mimics key processes engaged during mind-wandering and a metric describing a naturally occurring example of the state. No behavioural associations were recovered when using the DMN regions linked to autobiographical memory as a seed.

## Discussion

Our study set out to understand how the experience of mind-wandering while reading creates a state of perceptually-decoupled thought that derails our comprehension of the text. In particular, we tested an emerging hypothesis that spans both psychological and neural domains. At the psychological level, contemporary theories of self-generated states suggest that functional decoupling from perceptual input is important for effective memory retrieval, providing a process account of why mind-wandering during reading can derail comprehension (1). In the neural domain, we draw on a recent hypothesis that the role of DMN subnetworks in human cognition is related to their topographic location on the cortical mantle (18, 19). These regions occupy locations that are both the terminus of processing streams within the cortex important for abstract forms of cognition such as reading, and yet at a distance along the cortical surface from input systems, explaining why they can also be engaged by situations in which mental content is broadly unrelated to perceptual input (19).

Experiment 1 established a pattern of mutual inhibition between the act of reading for comprehension and the concurrent retrieval of autobiographically-relevant content. This pattern of mutual inhibition parallels the well-established negative correlation between naturally occurring mind-wandering and an individuals’ ability to comprehend what they are reading (7, 8). Using fMRI to index brain activity, we identified that these two states differentially recruited different aspects of the DMN, with greater recruitment of the dorsomedial DMN subnetwork when participants were reading for comprehension, and greater activity within the core of the DMN during autobiographical memory retrieval (See also 22). The involvement of the DMN in both self-generated mental content and reading for comprehension is consistent with prior studies exploring trait variation in mind-wandering while reading (16, 17).

Importantly, Experiment 1 found that ventral visual regions were engaged when participants were reading for comprehension as opposed to engaged in autobiographical memory recall, a pattern consistent with the view that the retrieval of autobiographical memories during mind-wandering creates a perceptually-decoupled state (1, 23). Experiment 2 used resting-state functional connectivity to establish that there is strong coupling at rest between regions of DMN relevant to reading and ventral visual cortex; in contrast, regions of core DMN, activated by autobiographical memory, showed reduced correlation with these ventral visual regions, compared with aspects of the DMN linked to reading. This pattern suggests that the reduction of activity in ventral visual cortex observed in Experiment 1 was not an artefact of our task instructions which required participants to attend to autobiographical memories rather than textual input. Instead, the core DMN regions activated when we retrieve autobiographical memories are functionally distant from regions of ventral visual cortex important for reading comprehension. Moreover, individual differences in intrinsic connectivity associated with the mind-wandering reading state are consistent with this view. We found that people who are more likely to mind-wander during reading show weaker coupling between reading-relevant DMN regions and an area of dorsal occipital cortex normally associated with autobiographical memory retrieval. In individuals who remain focussed on reading and who have better comprehension, this region of dorsal occipital cortex has stronger coupling with the aspects of DMN that are connected to visual regions that support reading.

Altogether our analyses have important implications for both psychological theories regarding self-generated states such as mind-wandering, as well as for our understanding of the involvement of the DMN in human cognition. Psychologically, our study suggests that simply asking individuals to retrieve autobiographical information creates a perceptually-decoupled state that is at odds with the comprehension of information from the external environment (1). This view assumes that perceptual decoupling is a condition related to the persistence of self-generated mental content in consciousness. In the neural domain, our data suggests that mind-wandering while reading involves a shift in the balance of neural activity within the DMN, away from lateral temporal and prefrontal regions that are closely linked to regions of ventral visual cortex important for reading, and towards regions within the core of this system involved in supporting mental content that is unrelated to the external environment during autobiographical memory retrieval. In this context, it is worth noting that studies that have established the role of the DMN in explicit memory retrieval also show that this pattern is accompanied by suppression of visual processing (e.g., 24). Furthermore, individuals with epilepsy, who are impaired in the process of pattern separation necessary for accurate episodic memory retrieval, show atypical suppression of brain activity in visual cortex during memory retrieval (25). The current data, therefore, provide novel support for the possibility that antagonistic activity patterns in the DMN and in sensory cortices may be important for features of memory retrieval to proceed in an effective manner.

More generally, our data add to growing evidence for a broad contribution of the DMN to features of human cognition. In particular, our findings are consistent with observations that the DMN can support apparently antagonistic states, particularly both perceptually-coupled and decoupled modes of cognition. Initial focus on the DMN assumed that this system was primarily important for internally-focused states. Recently, however, Yeshurun and colleagues (26) have argued that the DMN plays a key role in the integration of internal and external information in the service of aligning the perspectives of different individuals over time. Our data highlighting the role of the broader DMN in reading for comprehension is consistent with this perspective, as this mode of operation could help to create common ground between different individuals in their understanding of a narrative. However, the involvement of the same broad system in situations such as mind-wandering suggests that it can also lead to a breakdown in the common ground between individuals, in this case by impairing comprehension during reading. Critically, both of these different operational modes can stem from the DMN once it is recognised that the topographic location of this system at the end of sensory processing streams, such as the ventral visual stream, allows this network to provide abstract representations of external information. At the same time, this DMN location is functionally and structurally distant from unimodal sensory systems, providing the opportunity for decoupled states to emerge, which support mental contents that are unrelated to external input (such as self-generated off-task states) (18, 19).

Although our study provides important insight into how the occurrence of autobiographical mental content can derail our ability to make sense of the external environment, there are several important open questions that emerge from our study. First, in both Experiment 1, and in many examples of naturally occurring mind-wandering, the off-task mental contents may have greater relevance to the individual than information in the environment (14, 27), perhaps because these states rely on ventral regions of medial prefrontal cortex (11) that are important for motivated states (28–30). It is therefore unclear whether reading more engaging text would change the likelihood of off-task thoughts emerging, and the neural systems that are engaged during reading. To address this issue, it would be useful for future studies to explore the neural systems recruited by highly engaging and personally-relevant texts, to establish if these are more similar to those observed when reflecting on autobiographical memories. Second, our study established neural correlates that generalise across task-induced autobiographical memory (Experiment 1) and naturally occurring mind-wandering (Experiment 2), demonstrating important support for the process-occurrence view of self-generated states (1). However, our current data provide only limited constraints on how to understand the significance of the observed pattern of functional coupling. One possibility is that stronger coupling between the dorsomedial and core DMN subnetworks at rest is a neural fingerprint of individuals who are better able to maintain attention to what is being read (16, 17). It is also possible that for certain forms of comprehension, processing of visual information in dorsal occipital cortex is important, possibly depending on the white matter tracts identified through our supplementary analysis (See Supplementary Figure S6). Future research will be needed to establish why patterns of coupling between DMN subsystems are related to better comprehension of what is being read. Third, contemporary studies suggest that different types of self-generated thoughts have different neural correlates (31, 32). Accordingly, it is possible that certain features of naturally occurring off-task thought patterns could lead to greater or lesser disengagement from external input. This question could be readily addressed by combining multi-dimensional experience sampling (33) with brain activity recorded while individuals read.

## Methods

### Participants

A total of 339 participants were recruited in this study. For Experiment 1, 29 undergraduate students were recruited (age-range 18-23 years, mean age ± SD = 20.14 ± 1.26 years, 6 males). For Experiment 2, we used two separate resting-state samples: one sample consisted of 244 participants (age-range 18-31 years, mean age = 20.73 ± 2.39 years, 77 males; 3 participants overlapped with Experiment 1), which was used to examine the intrinsic connectivity of DMN regions identified in Experiment 1 that showed strong activity during reading and autobiographical memory retrieval. One participant was excluded from this analysis due to excessive head motion (i.e., mean head motion > .4 mm). Another dataset of 69 participants with behavioural measures of mind-wandering during reading (age range 18-31, mean age□=□19.87□±□2.33, 26 males; without any participant overlap with the other two samples) was used to characterise whether the functional architecture of the DMN regions associated with reading and autobiographical memory retrieval were related to individual variation in mind-wandering during reading, as measured in our prior study (17). All were right-handed native English speakers, and had normal or corrected-to-normal vision. None had any history of neurological impairment, diagnosis of learning difficulty or psychiatric illness. All provided written informed consent prior to taking part and received monetary/course credits compensation for their time. Ethical approval was obtained from the Research Ethics Committees of the Department of Psychology and York Neuroimaging Centre, University of York.

### Materials

144 highly imageable, frequent and concrete nouns were selected to serve as key words within sentences and as cue words for autobiographical memory recall. These nouns were divided into two lists (i.e., 72 words for each task) that did not differ in terms of frequency (CELEX database; 34), imageability (35), and concreteness (36; p > .1). The sentences were constructed by using these key words as a search term in Wikipedia to identify text that described largely unfamiliar facts about each item (Sentence Length: *Mean* ± *SD* = 20.04 ± .93 words). These sentences and the autobiographical memory cues were then divided into three sets and assigned to different conditions (with this assignment counterbalanced across participants). The sentences were assigned to (1) *Reading* without conflict from memory recall; (2) *Reading* with conflict from memory recall, and (3) *Memory Recall* with conflict from concurrent sentence presentation. Similarly, the autobiographical memory cues were assigned to (1) *Memory Recall* without conflict from sentences and (2) *Memory Recall* with conflict from sentences, as well as (3) *Reading* with conflict from memory recall. In addition, the words used in these conditions were matched on key psycholinguistic variables: they did not differ in lexical frequency, imageability, or concreteness (all *F* < 1.07). In addition, all the words in the three sets of sentences were comparable across these variables (see Table S1 in Supplementary material; all *F* < 1.40). Two additional cue words were created for task practice.

### Task Procedure for Reading and AM task

Testing occurred across two consecutive days. On Day 1, participants were asked to generate their own personal memories from cue words (i.e., Party) outside the scanner. They were asked to identify specific events that they were personally involved in and to provide as much detail about these events as they could, including when and where the event took place, who was involved, what happened, and the duration. They were asked to type these details into a spreadsheet, which ensured that comparable information was recorded for the different cue words.

On the following day, participants were asked to read sentences for comprehension, or to recall their generated personal memories inside the scanner. In reading trials, sentences were presented word by word, after either (1) an autobiographical memory cue word (e.g., Party), creating conflict between reading and personal memory retrieval, or (2) a letter string (e.g., XXX) allowing reading to take place in the absence of conflict from autobiographical memory. We controlled the duration of the sentences by presenting the words on 15 successive slides, combining short words on a single slide (e.g., *have been* or *far better)* or presenting articles and conjunctions together with nouns (e.g., *the need; and toys*). In memory recall trials, participants were asked to recall autobiographical memories during the presentation of either (1) an unrelated sentence, creating conflict from task-irrelevant patterns of semantic retrieval or (2) letter strings (XXX) allowing autobiographical memory to take place without distracting semantic input. As a control condition, meaningless letter strings (i.e., xxxxx) were presented. In order to ensure the participants were maintaining attention to the presented stimuli (even when these were irrelevant and creating competition), they were told to press a button when they noticed the colour of a word or letter string change to red. There were 3 trials out of 24 trials in each condition that involved responding in this way. Behavioural data for the colour change detection task are presented in Figure S1.

As shown in Figure 1, each trial started with a fixation cross (1–3s) in the centre of the screen. Then either an autobiographical memory cue word or a letter string appeared for 2s. During the presentation of the cue word, participants were asked to bring to mind their personal memories related to this item. Next, the task instruction (i.e., READING or MEMORY RECALL) was presented for 1s. Following that, words from sentences or letter strings were presented, with each one lasting 600ms. On memory recall trials, participants were asked to keep thinking about their autobiographical memory, in as much detail as possible, until the end of the trial.

After each trial, participants were asked to rate several dimensions of their experience. In the reading task, they were asked about task focus, their comprehension, and conceptual familiarity. For autobiographical memory trials, they were asked about task focus, vividness, and how consistent their retrieval was to the memory they specified day before. The three rating questions were sequentially presented after a jittered fixation interval lasting 1-3s. Participants were required to rate these characteristics on a scale of 1 (not at all) to 7 (very well) within 4s for each question. There were no ratings for the letter string trials.

Stimuli were presented in four runs, with each containing 30 trials: 6 trials in each of the four experimental conditions, and 6 letter string trials. Each run lasted 12.85 minutes, and trials were presented in a pseudorandom order to ensure trials from the same experimental condition were not consecutively presented more than three times.

Before entering the scanner, participants completed a 6-minute task to test their memory of the personal events they generated the day before scanning. They were also asked to review their generated memories and refresh themselves with the ones that were not well remembered. Next, they completed an 8-trial practice block containing all types of conditions to ensure fully understanding of the task requirements.

### Behavioural assessment of mind-wandering during reading

This dataset was used in our previous study (17). Participants were asked to complete a battery of behavioural assessments examining reading comprehension and off-task thought, while they read a passage about the topic of geology. During reading, they were required to note down any moments when they noticed they had stopped paying attention to the meaning of the text. After they finished reading, they were asked to answer 17 open-ended questions to assess their comprehension of the text, without being able to refer back to the text. A self-reported measurement, with 22 questions about the content of thoughts (e.g., *I thought about personal worries*), was used to assess off-task behaviour during the reading task (see Supplementary Figure S5). This analysis revealed that people were thinking about autobiographical memories (past events) alongside future events, other people and emotions, when they reported mind-wandering during reading. In this way, both off-task thoughts (i.e., frequency and the content of these experiences) and reading comprehension were assessed.

### Neuroimaging data acquisition

Structural and functional data were acquired using a 3T GE HDx Excite MRI scanner utilizing an eight-channel phased array head coil. Structural MRI acquisition in all participants was based on a T1-weighted 3D fast spoiled gradient echo sequence (repetition time (TR) = 7.8 s, echo time (TE) = minimum full, flip angle = 20°, matrix size = 256 × 256, 176 slices, voxel size = 1.13 × 1.13 × 1 mm^3^). The task-based activity was recorded using single-shot 2D gradient-echo-planar imaging sequence with TR = 3 s, TE = minimum full, flip angle = 90°, matrix size = 64 × 64, 60 slices, and voxel size = 3 × 3 × 3 mm^3^. In Experiment 1, the task was presented across 4 functional runs, with each containing 257 volumes. In Experiment 2, using the same scan parameters, a 9-minute resting-state fMRI scan was recorded, containing 180 volumes. The participants were instructed to focus on a fixation cross with their eyes open and to keep as still as possible, without thinking about anything in particular.

### Pre-processing of task-based fMRI data in Experiment 1

All functional and structural data were pre-processed using a standard pipeline and analysed via the FMRIB Software Library (FSL version 6.0, www.fmrib.ox.ac.uk/fsl). Individual T1-weighted structural brain images were extracted using FSL’s Brain Extraction Tool (BET). Structural images were linearly registered to the MNI152 template using FMRIB’s Linear Image Registration Tool (FLIRT). The first three volumes of each functional scan were removed in order to minimise the effects of magnetic saturation. The functional neuroimaging data were analysed using FSL’s FMRI Expert Analysis Tool (FEAT). We applied motion correction using MCFLIRT (37), slice-timing correction using Fourier space time-series phase-shifting (interleaved), spatial smoothing using a Gaussian kernel of FWHM 6 mm, and high-pass temporal filtering (sigma = 100 s) to remove temporal signal drift. In addition, motion scrubbing (using the fsl_motion_outliers tool) was applied to exclude volumes that exceeded a framewise displacement threshold of 0.9.

### Pre-processing of resting-state fMRI data in Experiment 2

Pre-processing was performed using the CONN-fMRI functional connectivity toolbox, Version 18a (http://www.nitrc.org/projects/conn; 38), based on Statistical Parametric Mapping 12 (http://www.fil.ion.ucl.ac.uk/spm/). Participants’ motion estimation and correction were then carried out through functional realignment and unwarping, and potential outlier scans were identified using the Artifact Detection Tool (ART) toolbox (https://www.nitrc.org/projects/artifact_detect). Structural images were segmented into Gray matter, White matter and Cerebrospinal Fluid tissues and normalized to the MNI space with the unified segmentation and normalization procedure (39). Functional volumes were slice-time (bottom-up, interleaved) and motion-corrected, skull-stripped and co-registered to the high-resolution structural image, spatially normalised to MNI space using the unified-segmentation algorithm (39), smoothed with an 8 mm FWHM Gaussian kernel.

Pre-processing steps automatically created three first-level covariates: a *realignment* covariate characterising the estimated subject motion for each participant, a *scrubbing* covariate containing the potential outliers scans for each participant, and a covariate containing quality assurance (QA) parameters (i.e., the global signal change from one scan to another and the framewise displacement) for each participant. Realignment parameters, potential outlier scans, signal from white matter and cerebrospinal fluid masks and effect of rest (i.e. an automatically estimated trend representing potential ramping effects in the BOLD timeseries at the beginning of the sessions), were entered as potential confound regressors into the model in the denoising step of the CONN toolbox. Using the implemented anatomical CompCor approach (40), all of these effects were removed in a single linear regression step to obtain a clean signal. Functional images were then band-passed filtered (.008 – .09 Hz) to constrain analyses to low-frequency fluctuations. A linear detrending term was also applied, eliminating the need for global signal normalisation (41,42).

### Analysis of task-based fMRI data in Experiment 1

The pre-processed time-series data were modelled using a general linear model, using FMRIB’s Improved Linear Model (FILM) correcting for local autocorrelation (43). Nine Explanatory Variables (EV) of interest and nine of no interest were modelled using a double-Gaussian hemodynamic response gamma function. The nine EVs of interest were: *Reading* (1) without and (2) with conflict from memory recall, *Autobiographical memory retrieval* (3) with and (4) without conflict from semantic input, (5) Letter String Baseline, (6–9) Task Focus effect for each of the four experimental conditions as a parametric regressor. Our EVs of no interest were: (10) Memory cue words and (11) Letter strings before the presentation of task instructions, Task instructions for *Reading* (12) without and (13) with conflict (this separation of the reading task instruction was based on the consideration that some recall or task preparation was likely to be occurring due to the presentation of autobiographical memory cues), plus task instructions for (14) *Memory Recall* and (15) *Letter String* baseline conditions. Other EVs of no interest were: (16) Fixation (the inter-stimulus fixations between the sentences or letter strings and the ratings questions), (17) Responses to catch trials (which included all time points with responses across conditions), and (18) Rating decision periods (including all the ratings across experimental conditions). EVs for each condition commenced at the onset of the first word of the sentence or the first letter string, with EV duration set as the presentation time (9s). The parametric EVs for the effect of Task Focus during the target had the same onset time and duration as the EVs corresponding to the four experimental trials, but in addition included the demeaned Task Focus ratings value as a weight. The fixation period between the trials provided the implicit baseline.

We examined the main effects of Task, and Conflict for both the main experimental conditions and the effect of Task Focus, and comparisons of each experimental condition with the letter string baseline, which allowed us to identify the activation and deactivation in each task. We report the results of each condition over letter baseline, the activation and deactivation of each task, and effects of task conflict in Supplementary Figure S2, S3, and S4. The four sequential runs were combined using fixed-effects analyses for each participant. In the group-level analysis, the combined contrasts were analysed using FMRIB’s Local Analysis of Mixed Effects (FLAME1), with automatic outlier de-weighting (44). A 50% probabilistic grey-matter mask was applied. Clusters were thresholded using Gaussian random-field theory, with a cluster-forming threshold of *z* = 3.1 and a familywise-error-corrected significance level of *p* < .05.

### Analysis of resting-state fMRI data in Experiment 2

The functional connectivity analysis was performed using DMN regions associated with reading and autobiographical memory retrieval as seeds. In a first-level analysis, we computed whole-brain seed-to-voxel correlations for each seed after the BOLD timeseries were pre-processed and denoised. For the group-level analysis of the dataset with 243 participants, we performed contrast between functional connectivity maps seeding from these two DMN seeds. For the group-level analysis of 69 participants, the EVs were entered into a GLM analysis, including reading comprehension scores, self-reported mind-wandering frequency, and the scores of the content of thoughts (for details see 17). We examined both the main effects and contrasted effects of these behavioural measures. In all analyses, we convolved the signal with a canonical haemodynamic response function. We used two-sided tests to determine significant clusters. Group-level analyses in CONN were cluster-size FWE corrected at *p* < .05, and used a height threshold of *p* < .005. Bonferroni correction was applied to account for the fact that we included two models, the *p*-value consequently accepted as significant was *p*□<□0.025. Prior to data analysis, all behavioural variables were z-transformed and outliers more than 2.5 standard deviations above or below the mean were imputed with the cut-off value. All brain figures were created using BrainNet Viewer (http://www.nitrc.org/projects/bnv/; 45).

### Neurosynth decoding

Task activation and conjunction maps were uploaded to Neurovault (46; https://neurovault.org/collections/9432/) and decoded using Neurosynth (47). Neurosynth is an automated meta-analysis tool that uses text-mining approaches to extract terms from neuroimaging articles that typically co-occur with specific peak coordinates of activation. It can be used to generate a set of terms frequently associated with a spatial map (as in Figure 2). The results of cognitive decoding were rendered as word clouds using free online word cloud generator (https://www.wordclouds.com/). We manually excluded terms referring to neuroanatomy (e.g., “inferior” or “sulcus”), as well as repeated terms (e.g., “semantic” and “semantics”).

## Author Contributions

M.Z., E.J., and J.S. designed research; M.Z., X.W., D.V., and K.K. performed research; M.Z., B.B., J.R., R.R., and R.V. analysed data; M.Z., E.J., D.S.M. and J.S. wrote the paper.

## Acknowledgments

This work was supported by the European Research Council (Project ID: 771863 – FLEXSEM to EJ), and a China Scholarship Council (CSC) Scholarship (No. 201704910952 to MZ).

