## Supporting information for "Perceptual coupling and decoupling of the default mode network during mind-wandering and reading"

***Supporting materials for Experiment 1***

***Linguistic properties of materials***

**Table S1.** *Linguistic properties of each set of key words within sentences and autobiographical memory cues, and the words within each set of sentences (M ± SD). Sets (i), (ii), and (iii), were counterbalanced across participants (see Materials).*

| ***Conditions*** | ***Frequency*** | ***Imageability*** | ***Concreteness*** |
| --- | --- | --- | --- |
| (i) sentence key words | 1.31 ± .56 | 591.67 ± 34.20 | 4.74 ± .52 |
| (ii) sentence key words | 1.47 ± .50 | 598.05 ± 27.49 | 4.72 ± .55 |
| (iii) sentence key words | 1.29 ± .52 | 592.61 ± 44.35 | 4.64 ± .58 |
| (i) autobiographical memory cues | 1.59 ± .76 | 588.48 ± 41.20 | 4.70 ± .47 |
| (ii) autobiographical memory cues | 1.48 ± .62 | 594.64 ± 24.46 | 4.75 ± .30 |
| (iii) autobiographical memory cues | 1.54 ± .56 | 601.53 ± 23.65 | 4.73 ± .41 |
| (i) sentence materials | 2.59 ± .24 | 354.34 ± 27.41 | 2.72 ± .26 |
| (ii) sentence materials | 2.52 ± .20 | 352.40 ± 23.39 | 2.72 ± .21 |
| (iii) sentence materials | 2.48 ± .27 | 347.64 ± 35.24 | 2.80 ± .24 |

***Behavioural results of catch trials***

**Results of catch trials**: Participants detected 75.6% of colour-change catch trials (i.e., they responded to this percentage of catch trials across conditions), showing that they were paying attention to inputs presented on the screen. Repeated-measures ANOVAs examining accuracy, RT, and response efficiency (i.e., RT divided by accuracy), and assessing the effects of Task (Reading vs. Autobiographical memory recall) and Conflict (No conflict vs. Conflict), revealed there were no differences in colour-change detection rates across conditions (see *Figure S1*); trials with no response were excluded from the RT analysis (24.4%). There was no main effect of Task (Accuracy: *F*(1,28) = 1.54, *p* = .22, *η_p_^2^* = .05; RT: *F*(1,28) = 1.92, *p* = .18, *η_p_^2^* = .06; Response efficiency: *F*(1,28) = .35, *p* = .56, *η_p_^2^* = .01), no main effect of Conflict (Accuracy: *F*(1,28) = .27, *p* = .61, *η_p_^2^* = .01; RT: *F*(1,28) = 2.83, *p* = .10, *η_p_^2^* = .001; Response efficiency: *F*(1,28) = 1.22, *p* = .28, *η_p_^2^* = .04), and no interaction (Accuracy: *F*(1,28) = .85, *p* = .36, *η_p_^2^* = .03; RT: *F*(1,28) = 1.27, *p* = .27, *η_p_^2^* = .04; Response efficiency: *F*(1,28) = .95, *p* = .34, *η_p_^2^* = .03).

We also performed paired-samples *t*-tests (Bonferroni-corrected for four comparisons) comparing colour-change detection for each experimental condition with the letter string baseline (RT: *M* ± *SD* = .53 ± .17 s; Accuracy: *M* ± *SD* = 74.7 ± 24.5 %; Response efficiency: *M* ± *SD* = .85 ± .57). Responses to baseline trials were significantly faster than for reading or recall trials (*t*(28) > 2.84, *p* < .009). For both accuracy and response efficiency, there were no significant differences between the experimental tasks and the letter string baseline data (*t*(28) < 1).

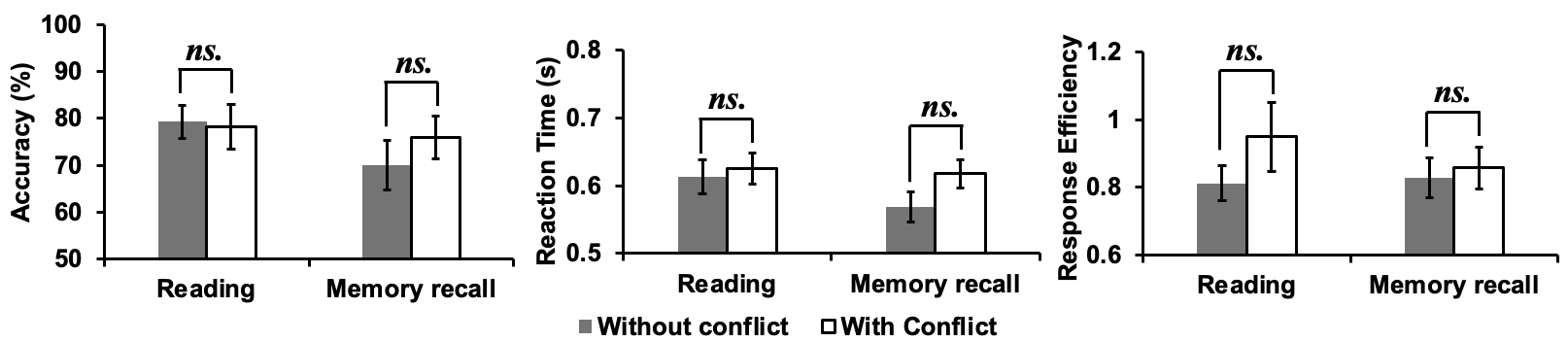

***Figure S1.* Behavioural results of catch trials.** Accuracy (percentage correct, left panel) and reaction time (in seconds, middle panel), as well as response efficiency (right panel) for the catch trials in each experimental condition (*Pure Reading*, *Conflict Reading*, *Pure Recall*, and *Conflict Recall*). Error bars represent the standard error. *ns.* indicates *not significant.*

***Effects of each experimental condition over letter string baseline***

For reading, the bilateral temporal regions (i.e., temporal poles, superior/middle/inferior temporal gyrus), precentral gyrus, middle/inferior frontal gyrus, temporal fusiform cortex, supplementary motor cortex, and visual cortex showed activation relative to the letter string baseline (see *Figure S2 A-B* of *Pure reading* > *Letter baseline* and *Conflict reading* > *Letter baseline* respectively). For autobiographical memory, middle temporal gyrus, temporal pole, middle/inferior frontal gyrus, insular cortex, supplementary motor cortex, and visual cortex showed activation compared to the letter string baseline (see *Figure S2 C-D* of the *Pure AM retrieval* > *Letter baseline* and *Conflict AM retrieval* > *Letter baseline*).

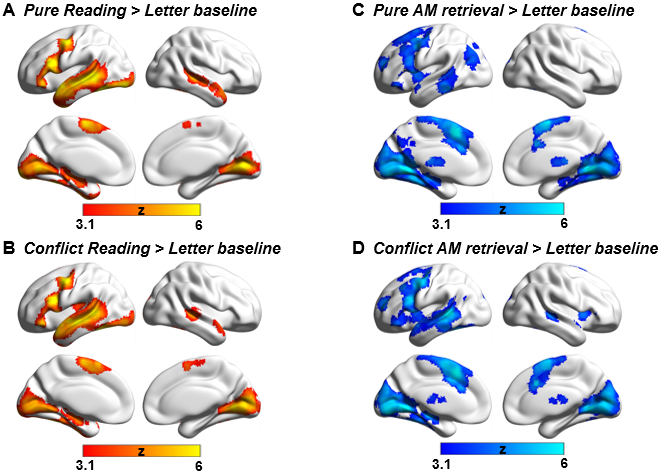

***Figure S2.* Comparisons between each experimental condition and meaningless letter string stimuli processing.** All maps were cluster-corrected with a voxel inclusion threshold of *z* > 3.1 and family-wise error rate using random field theory set at *p* < .05. L = Left hemisphere; R = Right hemisphere.

***Activation and deactivation in reading and autobiographical memory***

To identify activation and deactivation elicited by each task, we performed a formal conjunction analysis on the contrast maps of conflict and no-conflict for each experimental condition over the letter string baseline (providing a basic level of control for visual input and button presses). For reading, the bilateral temporal regions (i.e., temporal poles, superior/middle/inferior temporal gyrus), precentral gyrus, middle/inferior frontal gyrus, temporal fusiform cortex, supplementary motor cortex, and visual cortex showed activation relative to the letter string baseline (see *Figure S3 A*; the conjunction of *Pure reading* > *Baseline* and *Conflict reading* > *Baseline*), while bilateral middle frontal gyrus, supramarginal gyrus, medial prefrontal gyrus, anterior/posterior cingulate gyrus, and precuneus showed deactivation (see *Figure S3 B*; the conjunction of *Baseline* > *Pure reading* and *Baseline* > *Conflict reading*). For autobiographical memory, middle temporal gyrus, temporal pole, middle/inferior frontal gyrus, insular cortex, supplementary motor cortex, and visual cortex showed activation compared to the letter string baseline (see *Figure S3 C*; the conjunction of *Pure AM retrieval* > *Baseline* and *Conflict AM retrieval* > *Baseline*), while supramarginal gyrus showed deactivation (see *Figure S3 D*; the conjunction of *Baseline* > *Pure AM retrieval* and *Baseline* > *Conflict AM retrieval*).

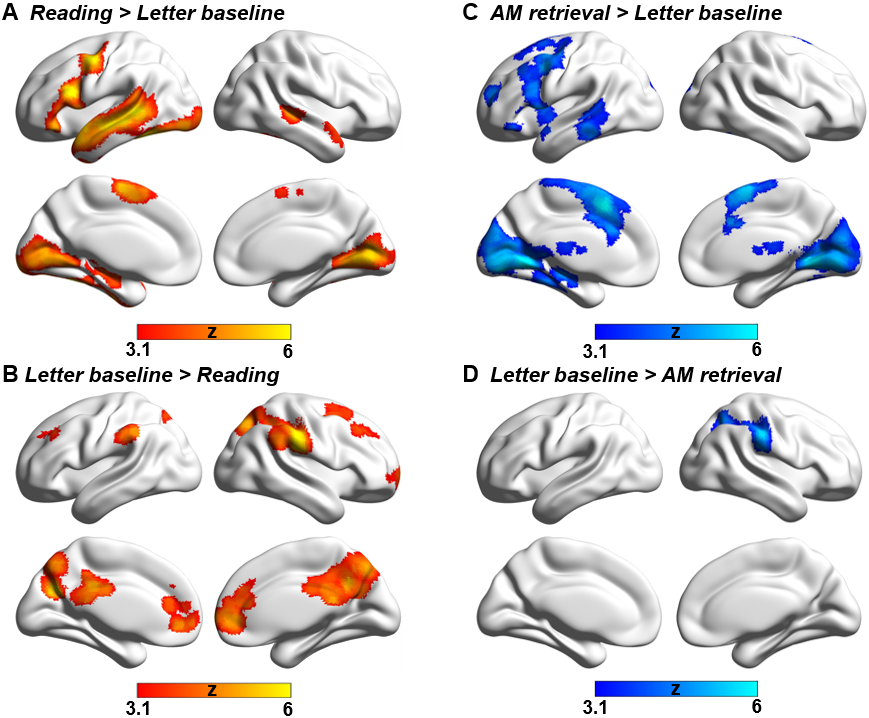

***Figure S3*. Task activation and deactivation. (A) *Reading* > *Letter baseline*** and **(B) *Letter baseline* > *Reading*** show the brain activation and deactivation during reading task relative to the letter string baseline. (**C) *AM retrieval* > *Letter baseline*** and **(D)** ***Letter baseline* > *AM retrieval*** show the brain activation and deactivation during autobiographical memory recall relative to the letter string baseline. These conjunctions were identified using FSL's ‘easythresh_conj’ tool. All maps were thresholded at *z* > 3.1 (*p* < .05). L = Left hemisphere; R = Right hemisphere.

***Effects of Task conflict***

We did not find any conflict effects for reading, over and above the task focus effects we present in the main manuscript. For autobiographical memory, the parahippocampus gyrus, temporal occipital fusiform, lateral occipital cortex, precuneus and anterior medial prefrontal cortex – identified as important areas for autobiographical memory by the *Autobiographical* *memory* > *Reading* contrast – showed greater activation when there was *no* distracting sentence input (see *Figure S4 A*). When there was conflict from irrelevant sentences during autobiographical memory recall, there was greater activation in some regions that were identified in the *Autobiographical* *memory* > *Reading* contrast, including precentral gyrus, frontal pole, frontal orbital cortex, and angular gyrus, plus greater activation in ventral visual cortex corresponding to the presentation of more complex visual input (see *Figure S4 B*).

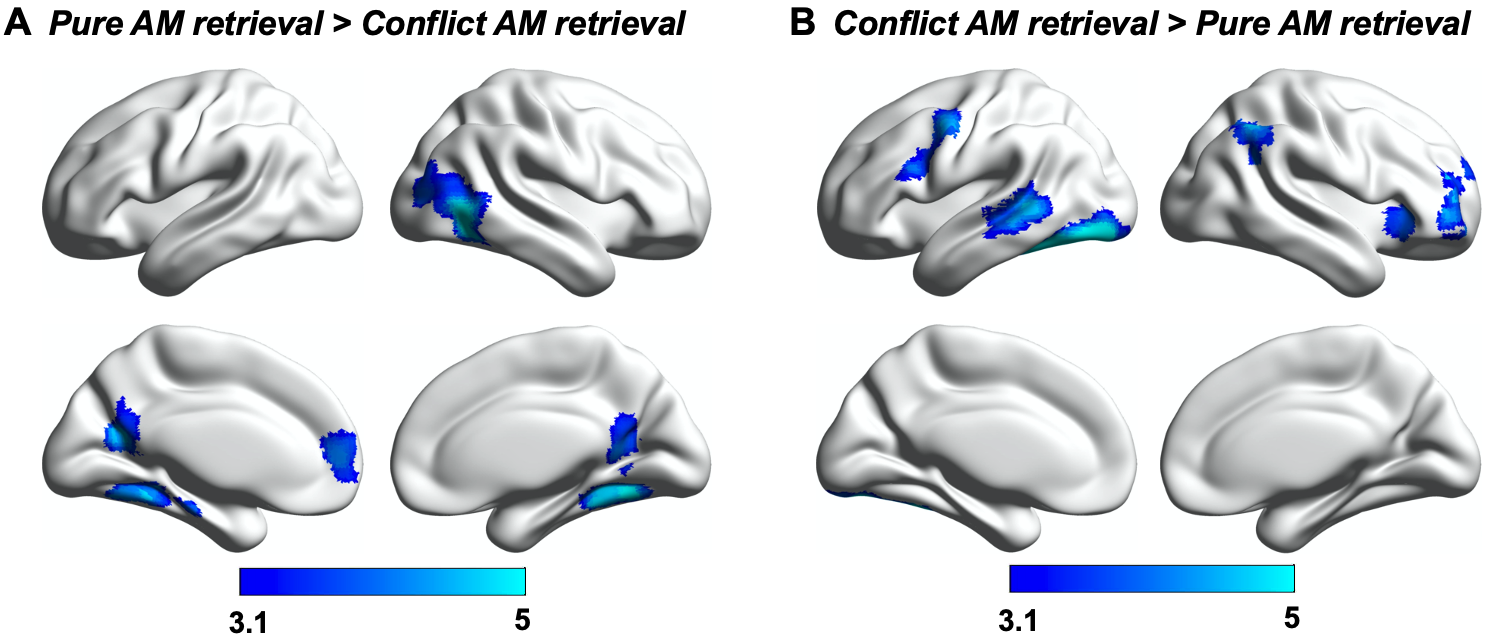

***Figure S4. Effects of Task conflict.* (A)** Significant activation when there was no conflict from semantic input defined using the contrast of *Pure AM retrieval* > *Conflict AM retrieval*. **(B)** Significant activation when there was conflict from semantic input defined using the contrast of *Conflict AM retrieval* > *Pure AM retrieval*. All maps were thresholded at *z* > 3.1 (*p* < .05).

***Supporting materials for Experiment 2***

***Contents of Mind-wandering while reading***

The assessment of mind-wandering behaviour contained 22 questions about the content of thoughts, rated on a scale of 1 (Completely did not describe my thoughts) to 9 (Completely did describe my thoughts). These questions refer to four types of mind-wandering contents, namely Past thoughts (e.g., *I thought about an event that took place earlier today*), Future thoughts (e.g., *I thought about an interaction I may possibly have in the future*), Social thoughts (e.g., *I thought about people I have just recently met*), and Emotional thoughts (e.g., *I thought about things I am currently worried about*). The descriptive rating for each type of mind-wandering content is shown in Figure *S5*.

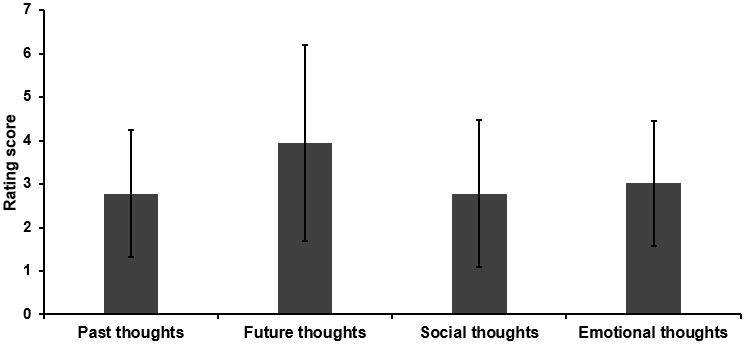

***Figure S5. Contents of mind-wandering during reading.*** Different types of internal thoughts that people reported that they were thinking about, when they mind-wandered during reading in Experiment 2.

***Structural connectivity analysis***

Our functional connectivity analysis revealed that reading-relevant DMN regions are more functionally connected to ventral visual cortex, compared to regions of DMN relevant to autobiographical memory. This effect might be due to the differences in the structural connectivity between visual cortex and these DMN subsystems. To examine this possibility, we performed white matter connectivity analysis using the visual network from Yeo et al. (1) as a seed, and reading- and AM-related DMN regions as masks, which allowed us to delineate patterns of structural connectivity from visual cortex to core and dorsomedial DMN subsystems.

To estimate structural connectivity between the visual network and reading- and AM-related DMN regions, we analyzed a pre-processed dataset of unrelated healthy young adults (N = 70, mean age = 31.97 ± 8.82 years old, 31 females). All processing was performed via micapipe, an openly accessible processing pipeline for multimodal MRI data (https://micapipe.readthedocs.io/). In brief, micapipe combines procedures from several software packages, including tools from AFNI, FSL, FreeSurfer, mrtrix, and ANTs. Cortical surfaces were generated by applying FreeSurfer to T1w scans (2), images were non-linearly aligned to MNI152 space using ANTs, and tissue types segmented using FSL FAST. The diffusion weighted imaging (DWI) data were pre-processed using MRtrix (3, 4). DWI data was denoised, underwent b0 intensity normalization, and were corrected for susceptibility distortion, head motion, and eddy currents using a reverse phase encoding from two b=0s/mm2 volumes. Required anatomical features for tractography processing (i.e., FSL-based tissue type segmentations) were co-registered to native DWI space using an affine transformation implemented in ANTs. Diffusion processing and tractography were performed in native DWI space. We performed anatomically-constrained tractography using tissue types (cortical and subcortical grey matter, white matter, cerebrospinal fluid) segmented from each participant’s pre-processed T1w images registered to native DWI space. We estimated multi-shell and multi-tissue response functions and performed constrained spherical-deconvolution and intensity normalization. To approximate ROI-to-ROI structural connectivity, we applied tckgen with a setting of 100,000 streamlines between the visual network and reading/AM components of the DMN in diffusion space, followed by generation of a tract-weighted image. Tract-weighted images were then mapped to MNI152 space using the existing co-registration/registation functions, and averaged across subjects to approximate consistent tracts across individuals. This analysis revealed that there are structural connectivity pathways from visual areas to DMN regions supporting reading and personal memory retrieval. The results are shown in *Figure S6*.

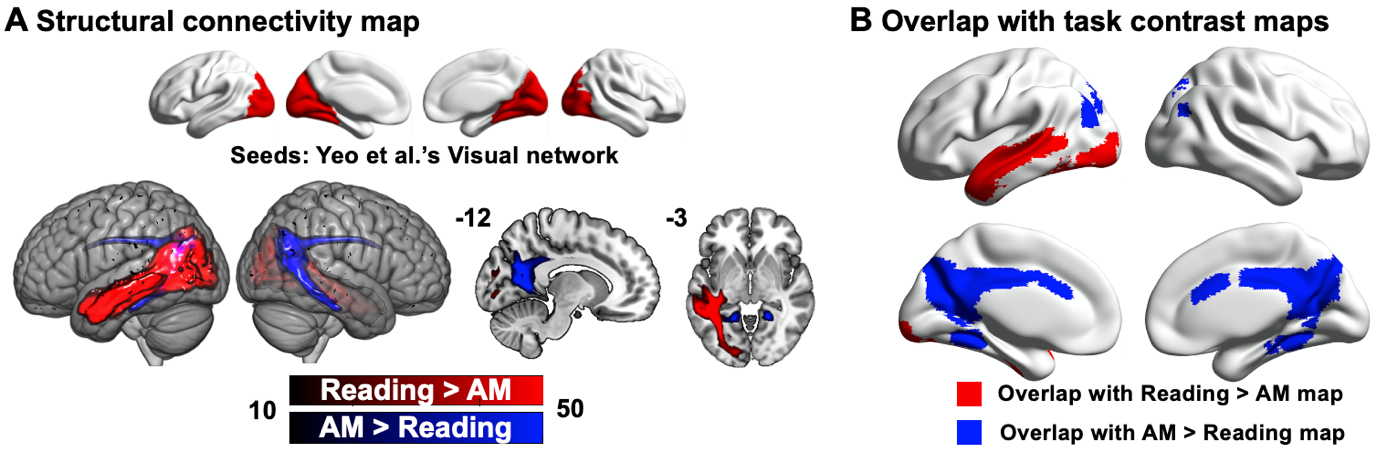

***Figure S6. Structural connectivity.* (A)** Structural connectivity seeding from visual network defined by Yeo and his colleagues (2011) to DMN regions linked to reading comprehension (in red) and autobiographical memory retrieval (in blue). **(B)** Overlap of structural connectivity map with *Reading > AM* and *AM > Reading* task contrast maps.

***Functional connectivity from reading-relevant visual cortex***

To further understand the role of dorsal occipital cortex (DOC) in reading comprehension (i.e., the cluster identified as having stronger decoupling from dorsomedial DMN in individuals more likely to mind-wander while reading), we performed resting-state functional connectivity analysis in a separate dataset (N = 243). This analysis explored how the functional connectivity of reading-relevant visual regions varies as a function of the connectivity strength between dorsomedial DMN and DOC (i.e., the connectivity pattern associated with real-world mind-wandering during reading in Experiment 2). First, we took reading-related DMN regions identified by the Reading > AM contrast (i.e., as shown in *Figure 2C*) as a seed and extracted the connectivity values for this seed within a mask defined by the DOC cluster in Figure 4C for each participant. Next, we seeded visual regions activated during reading, using the cluster defined by the contrast of Reading > AM from Experiment 1 as a seed. Finally, we searched for brain regions in which the intrinsic connectivity from this visual seed varied as a function of the connection between dorsomedial DMN and DOC. We found that for people with stronger connectivity between reading DMN and DOC, there was also stronger connectivity from the reading-relevant visual region to the same site within dorsal occipital cortex, along with supracalcarine cortex, lingual gyrus and precuneus cortex (see *Figure S7 A*). The overlap between this pattern of connectivity from reading-relevant visual cortex and the cluster associated with individual differences in mind-wandering while reading is shown in *Figure S7 B*. This finding suggests that the DOC region normally implicated in autobiographical memory retrieval (i.e. falling within task and connectivity contrast maps of AM > Reading, as well as the task focus effect for AM) is also more connected to ventral visual regions important for reading comprehension in people with stronger connectivity of this site to dorsomedial DMN. In this way, this DOC region may become more visually-coupled in people who have good reading comprehension and who avoid mind-wandering during reading.

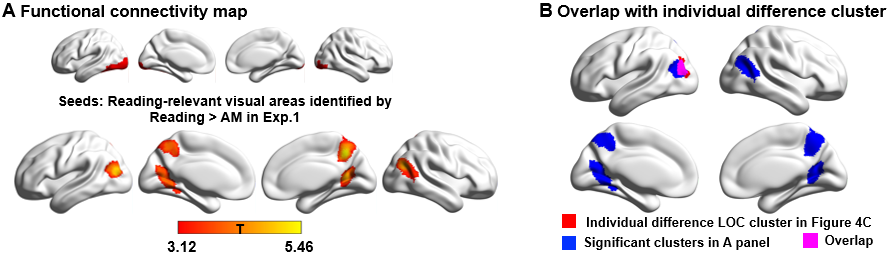

***Figure S7. Functional connectivity.* (A)** Functional connectivity seeding from reading-relevant visual areas identified by *Reading > AM* in Experiment 1. **(B)** Overlap with lateral occipital cortex (LOC) identified in individual difference analysis in Experiment 2.

***References in supplementary material***

1.	B. T. Yeo *et al.*, The organization of the human cerebral cortex estimated by intrinsic functional connectivity. *Journal of neurophysiology* **106**, 1125-1165 (2011).

2.	A. M. Dale, B. Fischl, M. I. Sereno, Cortical surface-based analysis: I. Segmentation and surface reconstruction. *Neuroimage* **9**, 179-194 (1999).

3.	J. D. Tournier, F. Calamante, A. Connelly, MRtrix: diffusion tractography in crossing fiber regions. *International journal of imaging systems and technology* **22**, 53-66 (2012).

4.	J.-D. Tournier *et al.*, MRtrix3: A fast, flexible and open software framework for medical image processing and visualisation. *Neuroimage* **202**, 116137 (2019).
